# Specific guidelines for time and space of multisensory plasticity in the superior colliculus

**DOI:** 10.1101/2023.02.12.528173

**Authors:** Linghong Wang, Yaxin Han, Zhe Sun, Biao Ouyang, Chao Dong

## Abstract

The potential to combine facts from exceptional senses, and thereby facilitate detecting and localizing events, commonly develops regularly in cat superior colliculus (SC) neurons. Multisensory integration in SC neurons depends on the spatial and temporal relationships of cross-modal cues. Here, we reveal that the parallel process of short-term plasticity during adulthood that would adapt multisensory integration to reliable changes in environmental conditions. The short-term experience altered the temporal preferences of SC neurons, this short-term plasticity was limited to changes in cross-modal timing (a factor commonly induced by events at different distances from the receiver. Nonetheless the plasticity was no evident in response to changes in cross-modal spatial configuration.

## Introduction

It is possible to describe the natural world as a never-ending stream of stimuli that change rapidly across the dimensions of space and time. The spatiotemporal complexity of these real-world stimuli continually challenges the nervous system’s ability to process information. As a result of evolution, brain structures have evolved that are designed specifically to process and integrate multisensory information. Among these structures, the superior colliculus (SC) is one of the best studied. It receives convergent input from the visual, auditory, and somatosensory systems. (Perrault et al. 2005; Wallace et al. 2004; Wang et al. 2020) Nevertheless, it appears necessary to synthesize information from systems that have fundamentally different operational parameters due to the extensive experience required to understand how to synthesize information. In mature multisensory superior colliculus (SC) neurons, sensory information is integrated across senses to enhance spatially and temporally congruent cross-modal responses.

Spatiotemporally concordant cross-modal cues typically elicit responses that are more robust than those to their component cues (multisensory enhancement). Cross-modal cues that are spatially or temporally disparate elicit less robust responses (multisensory depression).(Stein, Stanford, and Rowland 2014; Xu et al. 2012), these spatial and temporal principles guiding the products of integration spatially disparate configuration. are not inherent features of multisensory neurons. They are normally acquired through cross-modal experience during the first few months of early life.(Stein, Stanford, and Rowland 2009; Cuppini, Stein, and Rowland 2018)

This aim of the present study was to evaluate whether there was a parallel process of short-term plasticity during adulthood that would adapt multisensory integration to reliable changes in environmental conditions. If this is correct, it is necessary to Identify whether the presumptive plasticity of the system is equal across space and time.

## Materials and Methods

### Animals

Lab animals were cared for and used according to the Guide for the Care and Use of Laboratory Animals 8th Ed. (NRC 2011), and were approved by the Animal Care and Use Committee of Inner Mongolia Medical University.

### Surgical procedures

As anesthesia and tranquilization, ketamine hydrochloride was administered intramuscularly (25–30 mg/kg) and acepromazine maleate was given intravenously (0.01 mg/kg). In the surgical preparation room, the animal was given doses of prophylactic antibiotics (5 mg/kg enrofloxacin, intramuscularly) and analgesics (0.01 mg/kg buprenorphine, intramuscularly) and intubated. In the surgical suite, the animal was placed in a stereotaxic frame and deep anesthesia was induced and maintained (1.5–3.0% inhaled isoflurane). During the surgery, expired CO2, oxygen saturation, blood pressure, and heart rate were monitored with a vital signs monitor (Philips EarlyVue VS30) and body temperature was maintained with a hot water heating pad. Stainless steel screws and dental acrylic were used to secure a stainless-steel recording cylinder to the skull following a craniotomy dorsal to the SC. After suturing the skin around the implant, the anesthetic was discontinued and the animal was allowed to recover. After the animal regained mobility, analgesics were administered in its home pen. (2 mg/kg ketoprofen, IM, once per day; 0.01 mg/kg buprenorphine, IM, twice per day) for up to 3 d.

### Recording procedures

A minimum of seven days after recovery, the animal began weekly recording sessions. Each recording session involved anesthesia and tranquilization with ketamine hydrochloride (20 mg/kg, IM), as well as acepromazine maleate (0.1 mg/kg IM), Intubated, artificially respired, and secured to a stereotaxic frame in a recumbent position by attaching two head posts to a recording chamber, which ensured no wounds or pressure points. A constant end-tidal CO2 level of 4% was achieved by adjusting the respiration rate and volume. Expiration CO2, pulse rate, and blood pressure were monitored continuously to assess metabolic status, if necessary, adjust the depth of anesthesia. In order to avoid movement artifacts and fix the orientation of the eyes and ears, rocuronium bromide injections (0.7 mg/kg, IV) were administered to induce neuromuscular blockade. A contact lens was placed over the eyes so that the cornea would not dry out and visual stimulus could be focused on the tangent plane. During the procedure, ketamine hydrochloride (5 mg/kg/h), rocuronium bromide (1–3 mg/kg/h) 5% dextrose solution (2–4 ml/h), and saline were infused intravenously to maintain anesthesia, neuromuscular blockade, and hydration. The body temperature was maintained using a hot water pad at 37–38°C.

To locate single neurons in the SC’s multisensory layers, an electrode manipulator used to lower the glass-coated tungsten electrode was advanced by hydraulic Microdrive. The neural data were sampled at ~24 kHz, band-pass filtered between 300 and 3000 Hz, and spike-sorted online and offline using a Jiangsu Brain Medical Technology recording system and sorter software. When a visual–auditory neuron was isolated (amplitude 4 or more SDs above background), its visual and auditory receptive fields (RFs) were mapped manually using moving light bars and broadband noise bursts. These stimuli were generated from a projection screen and speakers ~30 cm from the animal’s head.

### Test Stimuli

Visual stimuli have been shifting mild bars (size: 6° × 20°; intensity: 5–35cd/m^2^) and flashing stationary spots (size: 4°; duration: 50 ms; intensity: 4.8–13.1 cd/m^2^). Auditory stimuli have been 100-ms broadband noise bursts of 60–75 dB SPL, delivered with the aid of audio system established on a 30-cm hoop and separated with the aid of 15°. The hoop should be circled round the head.

For every neuron, visual, and auditory receptive fields have been mapped with the visible and auditory check stimuli and a take a look at region used to be chosen inside a rather responsive region of its overlapping receptive fields in order to grant the best possible chance of exposing multisensory integration (Kadunce et al. 2001) (Ghose and Wallace 2014)Then, visible and auditory check stimuli have been introduced in my view and in spatiotemporally concordant combos in pseudo-random order. To similarly decrease the opportunity of erroneously concluding that a neuron used to be incapable of multisensory integration as an end result of cortical deactivation, every neuron that failed to show off statistically big response enhancement to the preferred multisensory take a look at battery was once subjected to extra tests. These concerned stimuli at a couple of stimulus onset asynchronies and a couple of places inside the vicinity of receptive discipline overlap, and three distinct stimulus parameters (intensity, motion direction, bar lengths, or stationary flashed spots).

### Stimulus configuration

This was examined in visual-auditory neurons using a bar of light moved through the receptive field and a broadband noise burst. Once per week adult animals were repeatedly presented with many (100) exposure trials of these visual-auditory (VA) cues in one of 3 configurations:

a. Spatiotemporally concordant (V leading A by 50ms),
b. Spatially disparate (V inside & A outside its receptive field),
c. Temporally disparate (V leading A by 100ms or 150ms).

Figure1 shown are the various exposure (“Training”) configurations presented on schematics of the visual and auditory fields (each circle=10°). Left: The stimuli were in close spatiotemporal concordance (“V” and “A” at 50ms SOA - generally the optimal Stimulus Onset Asynchrony). Middle: In the spatially disparate condition the stimuli were temporally coincident, but the auditory stimulus was outside the neuron’s excitatory receptive field. Right: The two spatial configurations were used to examine the effect of temporal disparity. In this case each configuration was presented with the visual stimulus preceding the auditory by 100 or 150ms.

**Figure1.**
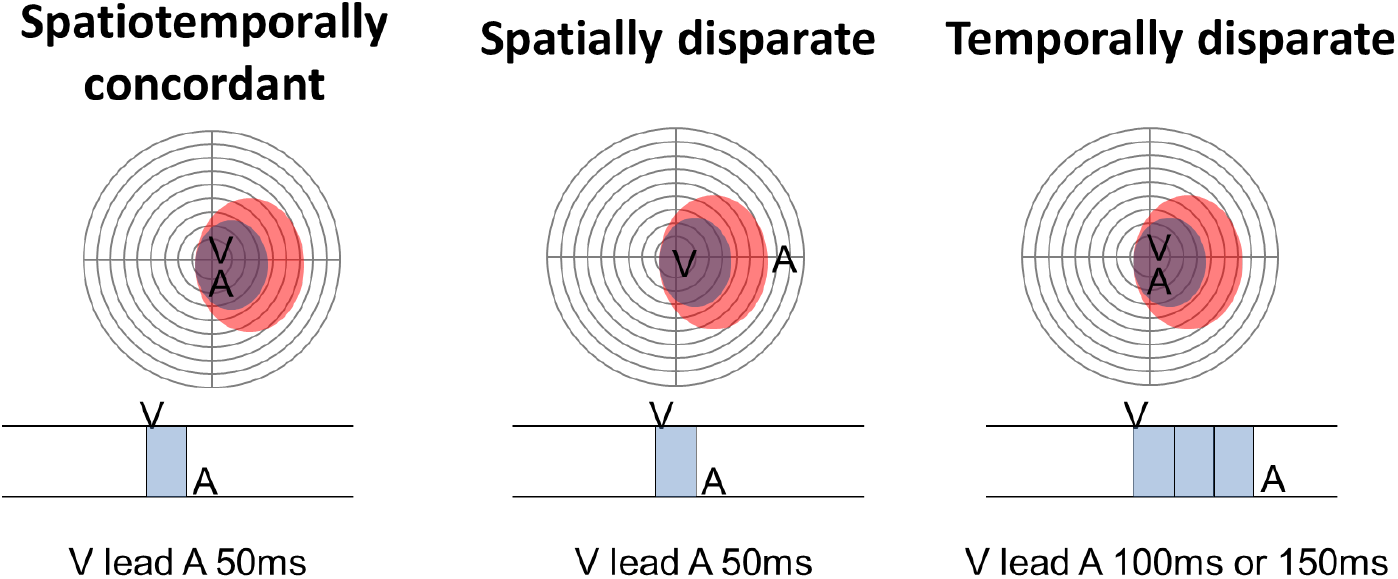
visual-auditory stimulus configuration

### Data analysis

Impulse instances (1 ms resolution) have been recorded for every trial and analyzed off-line. The response window was once described the usage of the algorithm recognized in beforehand studies.(Rowland and Stein 2008; Wang et al. 2020) The magnitude of every response used to be recognized as the imply range of impulses happening in the response window minus the anticipated range given the spontaneous firing charge (mean spontaneous firing price for every situation was once calculated in the 500ms window previous the stimulus). The response to the blended cross-modal stimuli used to be statistically in contrast with the response evoked via the extra tremendous person issue (i.e., modality-specific) stimulus to decide if the neuron’s response used to be considerably stronger (t test, alpha = 0.05). Multisensory enhancement used to be used as the index of multisensory integration, as it has been proven to be the most touchy index of an SC neuron’s multisensory integration capabilities(Kadunce et al. 2001). Multisensory enhancement (ME) was quantified by using percentage distinction between the response to the cross-modal stimulus and the response to the most positive modality-specific element stimulus. Statistical tests included the t-test and Mann–Whitney U, ANOVA, and chi-square tests as appropriate (alpha = 0.05). Summary statistics were expressed as mean ± standard error.

## Result

The essential findings of the experimental sequence are summarized by using the exemplars supplied in Figure.2, Figure.3 and Figure.4 Exemplar neurons illustrating the results of cross-modal testing in various conditions.

**Figure 2:**
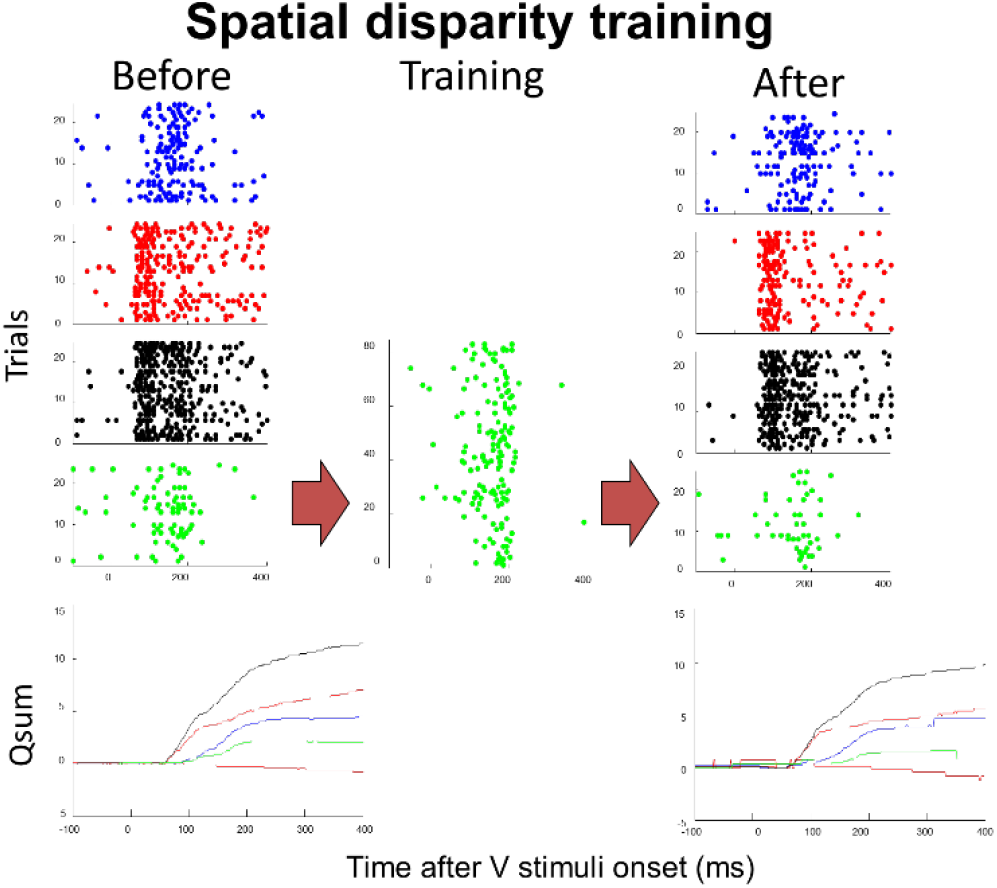
Spatial disparity training.

**Figure 3:**
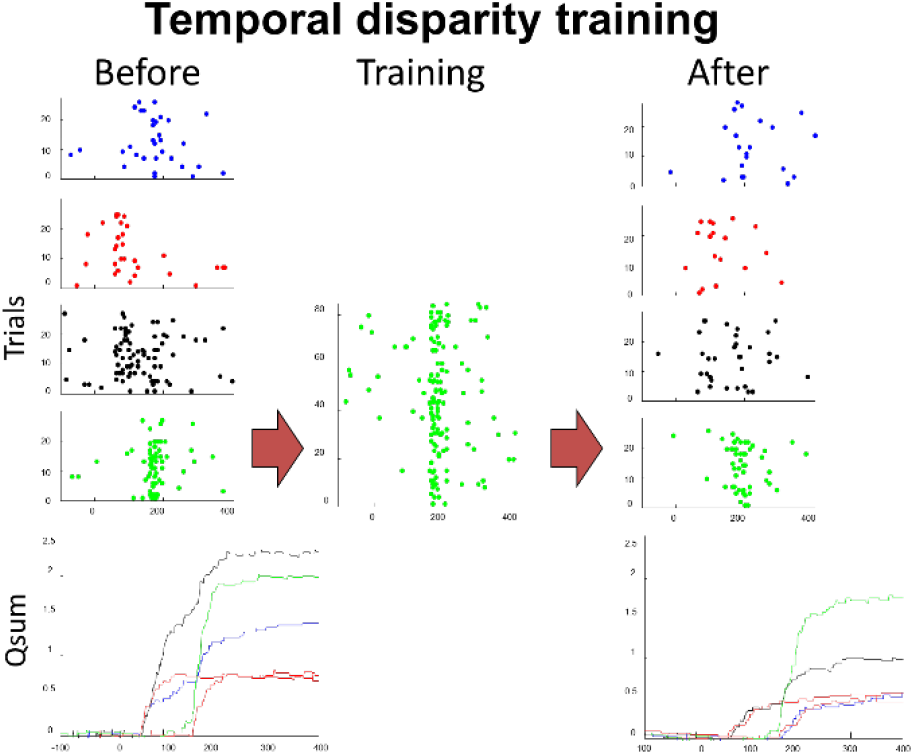
Temporal disparity training neuron type 1

**Figure 4:**
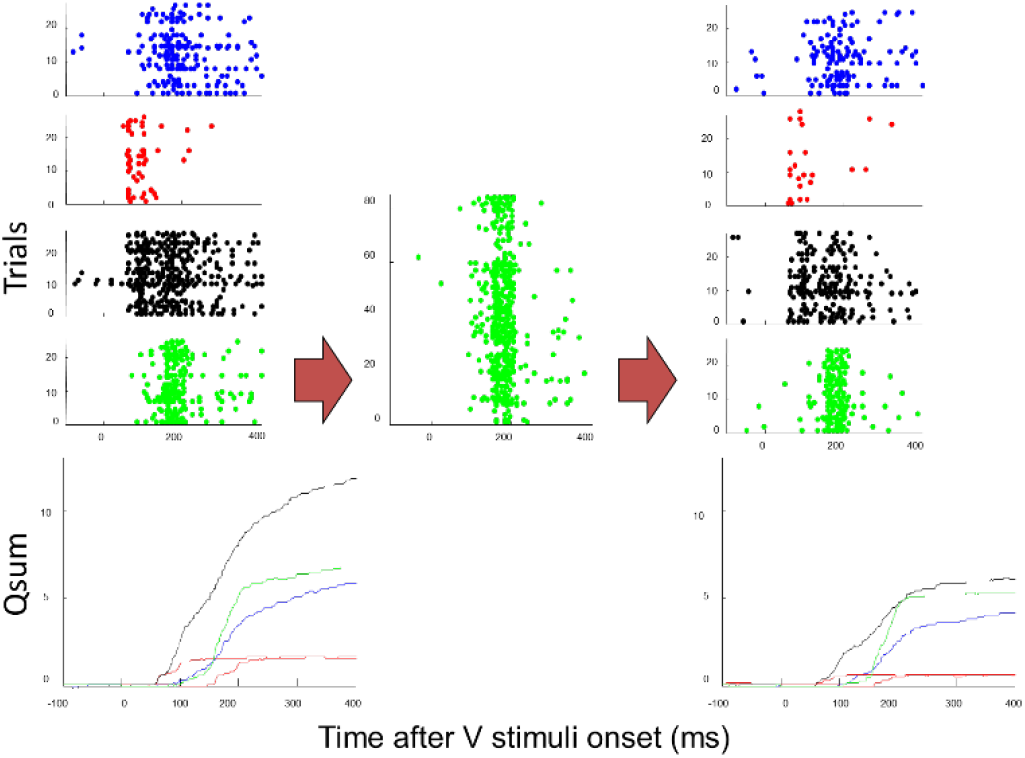
Temporal disparity training neuron type 2

Rasters show responses of this exemplar neuron before repeated exposure to the “Training” stimulus. From top-bottom the series of rasters show: Its unisensory visual responses (blue), auditory responses (red), multisensory responses to spatially coincident cross-modal stimuli (black) and multisensory responses to disparate cross-modal stimuli (green). Center: Responses to the disparate cross-modal training configuration. Right: The same sequence of responses is shown after training. Note the absence of obvious changes in the relative efficacy of the various stimulus configurations. Cumulative impulse counts (QSum) for each response are displayed at the bottom using the same color conventions as above. The response to the spatiotemporally concordant V-A cues (black) is significantly (p<0.05) more robust than the most effective unisensory response (auditory) both before and after training. The response to the spatially disparate cross-modal configuration (green) was significantly (p<0.05) less robust than the visual response both before and after training. However, none of the response magnitudes changed significantly before and after training.

Temporal disparity training can reverse a neuron’s SOA preference: The conventions here are the same as in Figure 1, except the rasters and qsums plotted in green now represent the responses to spatially concordant but temporally disparate visual and auditory cues (SOA=150ms). Prior to training with this temporally disparate configuration, the neuron exhibits significantly (p<0.05) enhanced responses to the spatiotemporally concordant and temporally disparate cross-modal configurations. There is a slight (not significant) preference for the spatiotemporally concordant configuration. However, repeated exposure to the temporally disparate configuration causes a significant (p<0.05) reduction in the neuron’s response to the spatiotemporally concordant cross-modal configuration. After training, the neuron’s responses to concordant and temporally disparate cross-modal configurations are still enhanced over the unisensory responses (p<0.05); however, the response to the temporally disparate configuration is now significantly more robust than the response to the temporally concordant configuration (p<0.05). This reversal of SOA preference was noted in 60% neurons.

Temporal disparity training can flatten a neuron’s SOA preferences: Conventions here are the same as in Fig. 3. In this case the neuron’s preferences were not reverse but substantially attenuated, thereby flattening its SOA tuning function. Left: The response of the neuron to spatiotemporally concordant cross-modal cues is significantly enhanced over the most robust (visual) unisensory response (p<0.05); however, the response to the temporally disparate configuration (SOA=150 ms) is not initially significantly enhanced. There is a significant difference (p<0.05) between the two multisensory responses, indicating a preferential tuning for the concordant configuration. Right: this preference was markedly diminished after training, so that the difference between the responses to the concordant vs. disparate configuration was now non-significant. As in Fig. 3, this change was association with a significant decrease (p<0.05) in the response to the concordant configuration.

### Population results of spatial disparity training

(see Figure 5.) Left: Shown are the mean changes in responses (impulses) as a consequence of training for each stimulus type. Note the general decrease in the unisensory responses (blue, red), but no change in the multisensory response to the spatiotemporal coincident configuration (VA50). Right: This lack of training effect was also evident when evaluating changes in the magnitude of the multisensory response (ME). Before training the coincident stimulus configuration produced a multisensory enhancement index of 55%, whereas the disparate configuration produced an index of –25%. Training did not significantly change either value (ns = non-significant).

**Figure 5.**
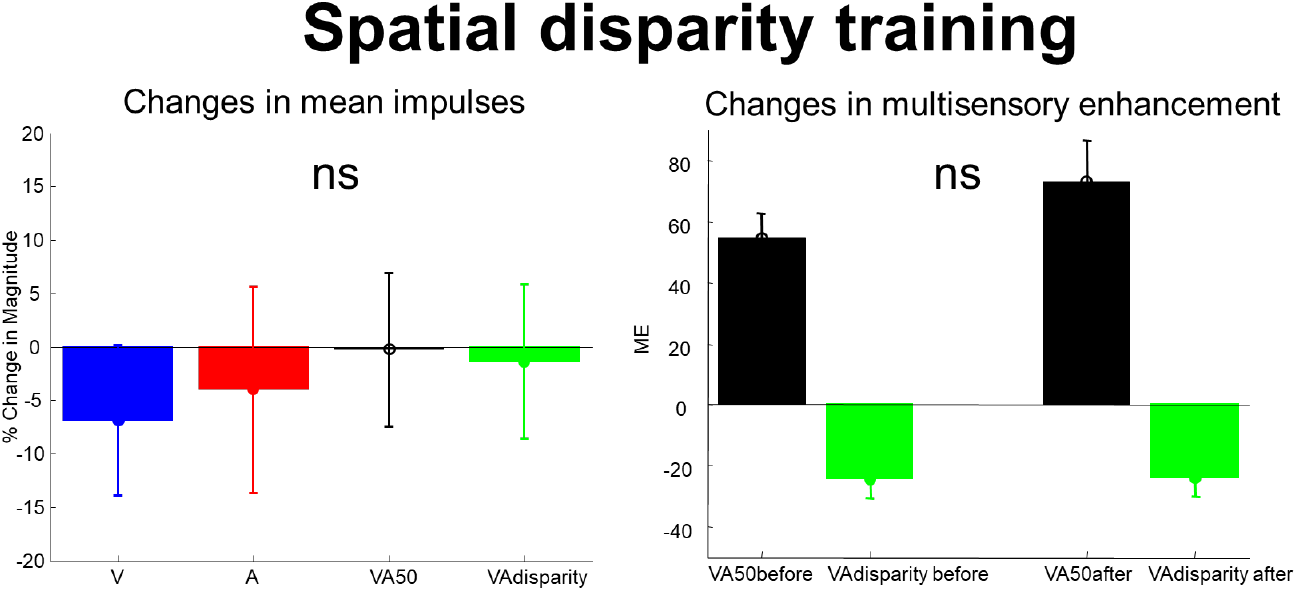
Population results of spatial disparity training

### Population results of temporal disparity training at 150ms SOA

(see Figure 6.) Conventions are the same as in Figure 5. Left: Note that training with an SOA of 150ms had negligible effects on the mean number of impulses in the unisensory responses. However the original preferred SOA of 50ms now elicited far fewer impulses whereas the training SOA of 150ms now elicited significantly more impulses. Thus, neuron’s changed their response profiles toward the training SOA. Right: This change in preference was also evident in the magnitude of the multisensory enhancement index. The ME to the 50ms SOA now decreased significantly, and the ME to the 150ms SOA increased significantly. * = p<.05.

**Figure 6.**
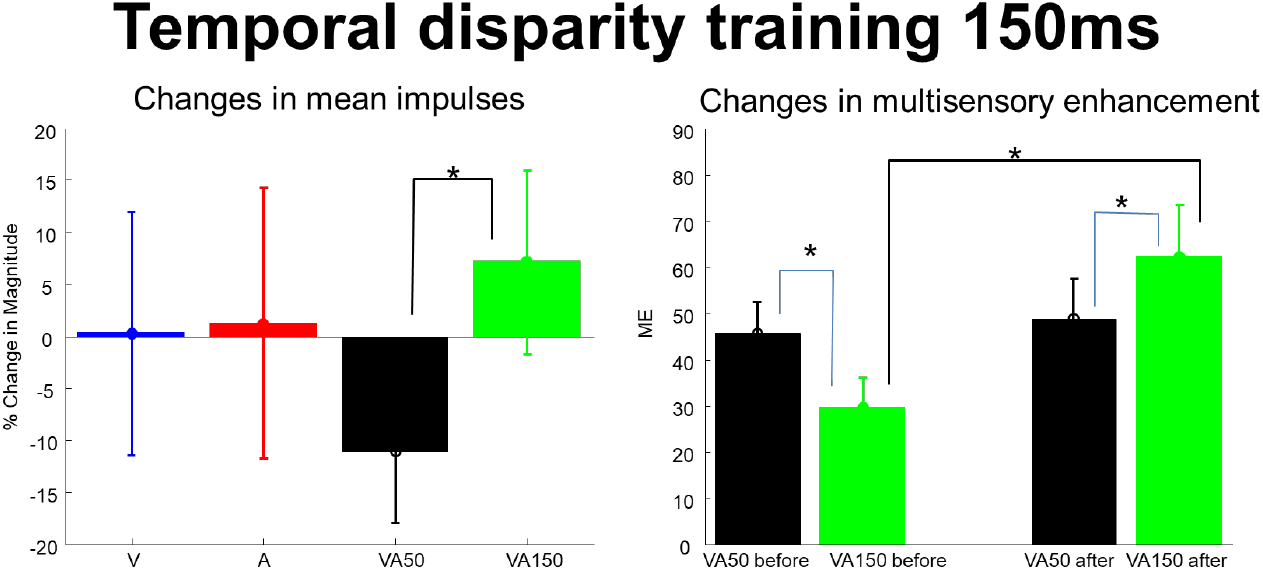
Population results of temporal disparity training at 150ms SOA.

### Population results of temporal disparity training at 100ms SOA

(see Figure 7.) In contrast to the significant changes induced by training with an SOA of 150ms, training at an SOA of 100ms induced no reliable response changes, although the trend is in the same direction. This marginal trend may be due to the fact that neurons show less preferential tuning between SOAs of 50ms and 100ms as they do between SOAs of 50 and 150ms.

**Figure 7.**
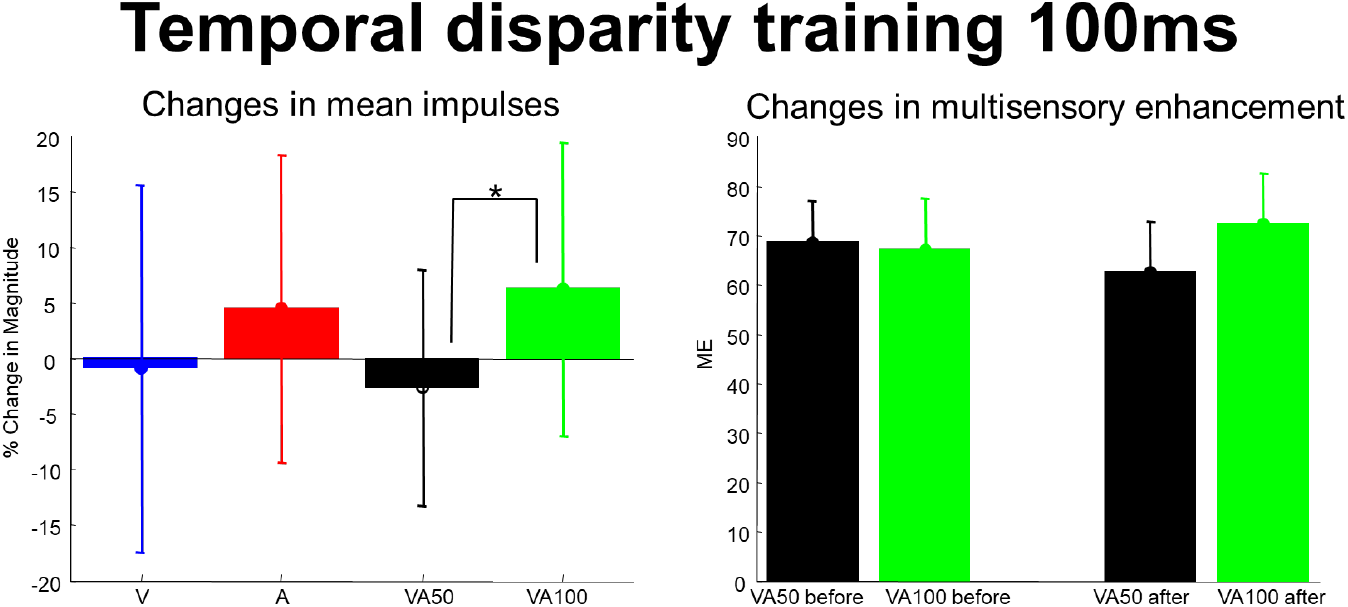
Population results of temporal disparity training at 100ms SOA.

## Discussion

The present study demonstrated that short-term experience altered the temporal preferences of many neurons, sometimes switching their temporal preferences toward the SOA presented in the exposure trials, and in others reducing their original preference. In contrast, there was little effect on their spatial preferences-regardless of whether the exposure was to spatially concordant or spatially disparate configuration.

Adult SC neurons retain the ability modify multisensory responses based on experience.(Xu et al. 2015). However, there are few experimental data on short-term plasticity in superior colliculus multisensory neurons. Our data shows that short-term plasticity was limited to changes in cross-modal timing (a factor commonly induced by events at different distances from the receiver. Nonetheless the plasticity was no evident in response to changes in cross-modal spatial configuration.

In sum, the current observations divulge the momentary experience altered the temporal preferences of some superior colliculus multisensory neurons. Whereas, the short-term plasticity used to be response to adjustments in cross-modal spatial configuration remains to be determined.

## Supporting information

References

